# Deep learning-based predictions of gene perturbation effects do not yet outperform simple linear baselines

**DOI:** 10.1101/2024.09.16.613342

**Authors:** Constantin Ahlmann-Eltze, Wolfgang Huber, Simon Anders

## Abstract

Advanced deep-learning methods, such as foundation models, promise to learn representations of biology that can be employed to predict *in silico* the outcome of unseen experiments, such as the effect of genetic perturbations on the transcriptomes of human cells. To see whether current models already reach this goal, we benchmarked five foundation models and two other deep learning models against deliberately simplistic linear baselines. For combinatorial perturbations of two genes for which only the individual single perturbations had been seen, we find that the deep learning-based approaches did not perform better than a simple additive model. For perturbations of genes that had not yet been seen, the deep learning-based approaches did not outperform the baseline of predicting the mean across the training perturbations. We hypothesize that the poor performance is partially because the pre-training data is observational; we show that a simple linear model reliably outperforms all other models when pre-trained on another perturbation dataset. While the promise of deep neural networks for the representation of biological systems and prediction of experimental outcomes is plausible, our work highlights the need for clear setting of objectives and for critical benchmarking to direct research efforts.

---

The success of large language models in knowledge representation has spawned research efforts to apply foundation models to biology.^1,2,3^

The expectation is that such models can ingest enormous amounts of biological data and learn enough about biological systems, with their gene regulatory networks, metabolic pathways, protein complexes, and cellular architecture, to then reliably *generate* the outcome of unseen experiments—in analogy to generative models that produce meaningful texts or images upon a user’s prompt. Such capabilities could partially replace expensive experiments and enable exploration of otherwise intractable experimental designs. Datasets that are believed to have the appropriate scale are becoming available, such as the Human Cell Atlas,^4^ and are collected by projects such as CELLxGENE,^5^ which hosts millions of single-cell gene expression profiles from across species, organs, health and disease states.

Several single-cell foundation models have been published in the last three years.^6,7,8^ Trained on data from millions of single cells, they are intended to be useful for various downstream tasks. The pre-trained models can be downloaded, and a user can apply them directly to specific tasks or further fine-tune them with additional data.

Two recent foundation models, scGPT^9^ and scFoundation,^10^ promise to be able to predict the effect of genetic perturbations on the transcriptome. Their creators compared their performance against GEARS^11^ and CPA,^12^ two other deep learning models that tackle this task using graph neural networks and variational autoencoders, respectively. In this study, we benchmark the performance of these models in predicting gene expression changes after genetic perturbation against each other and against deliberately simple baselines. In addition, we consider three more single-cell foundation models: scBERT,^6^ Geneformer^7^ and UCE.^8^ While these were not explicitly designed to predict transcriptomic changes from *in silico* perturbations, they can be made to do so by combining them with a linear decoder that maps the cell embedding to the gene expression space. In the figures, we mark them with an asterisk in order to acknowledge that for these tools, in contrast to the former ones, no claim of suitability for the prediction task has been made.

## Results

### Predicting the effect of double perturbations

We used the dataset of Norman et al.^13^, where the expression of 100 individual genes and 124 pairs of genes in K562 cells was, each in turn, upregulated with a CRISPR-based activation system (overview in Suppl. Fig. S1). The phenotypes for these 224 perturbations plus the control condition without perturbation are the logarithm-transformed RNAseq expression values for 19 264 genes, which we downloaded from the scFoundation publication (see Data Availability). We fine-tuned the models on all 100 single and 62 of the double perturbations and assessed the prediction error on the remaining 62 double perturbations. We ran each analysis five times using different random partitions of the 124 double perturbations into two halves.

For the purpose of benchmarking, we included two simple baselines: (1) the *no change* model that always predicts the same expression as in the control condition, and (2) the *additive* model that, for each double perturbation, predicts the sum of the individual log fold changes (LFC) compared to the control condition. Neither uses any of the double perturbation data.

We measured each model’s prediction error as the *L*_2_ distance between predicted and observed expression values for the top 1 000 most highly expressed genes. To increase the robustness of our conclusions, we also examined further summary statistics, such as the Pearson delta measure, and other gene subsets: the *n* most highly expressed or *n* most differentially expressed genes, for various *n*. Focusing the assessment on such *read-out* gene subsets rather than the full set of 19 264 genes is appropriate because the data for the remaining majority of genes is more strongly affected by detection limit and noise. We found that all of the models had a prediction error substantially higher than the additive baseline (Figure 1A, B; Suppl. Fig. S2).

**Figure 1.**
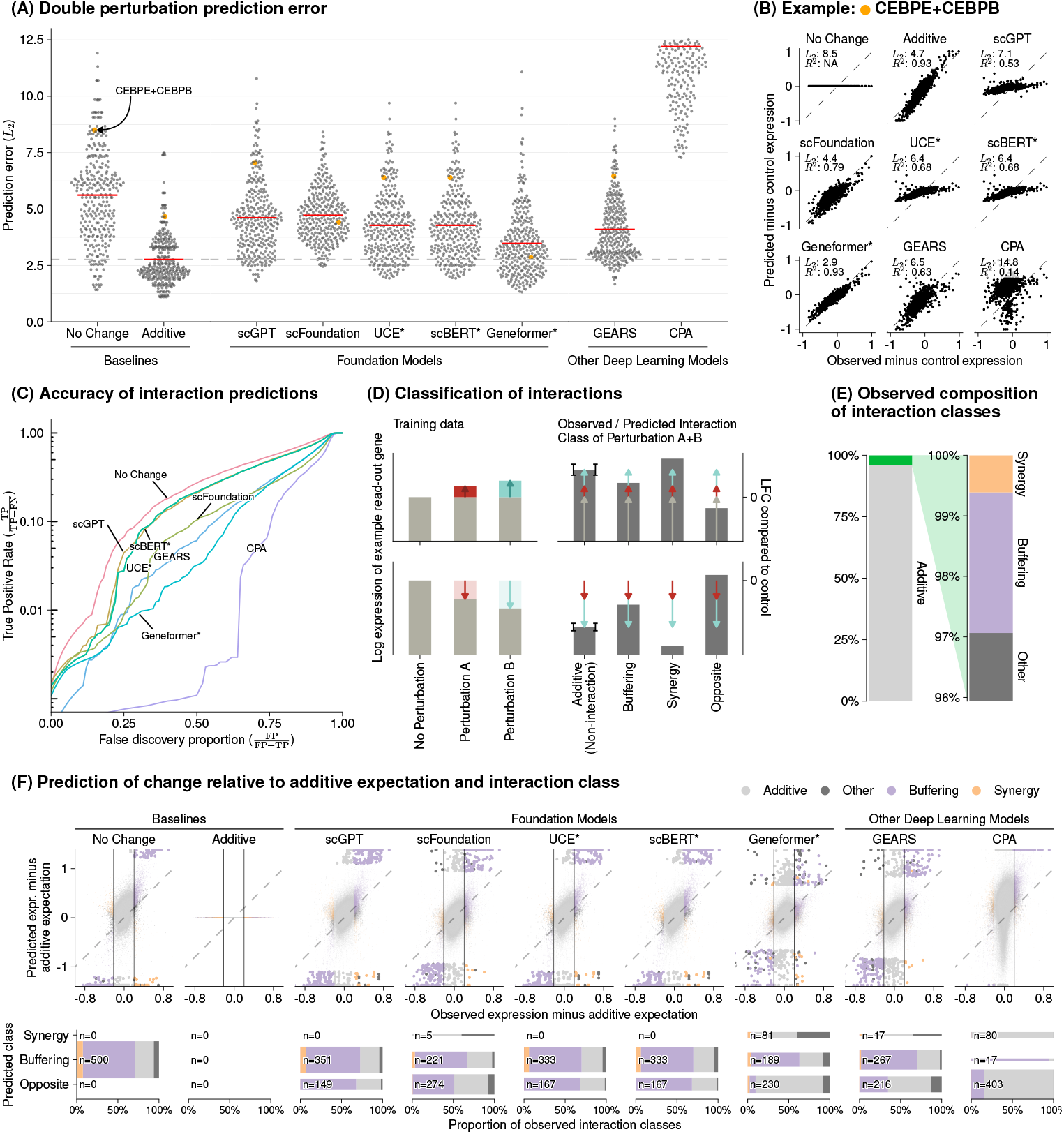
Double perturbation prediction. (A) Beeswarm plot of the prediction errors for 62 double perturbations across five test-training splits. The prediction error is measured by the *L*_2_ distance between the predicted and observed expression profile of the *n* = 1000 most highly expressed genes. The horizontal red lines show the mean per model, which for the best-performing model is extended by the dashed line. (B) Scatterplots of observed vs. predicted expression from one example of the 62 double perturbations. The numbers indicate error measured by the *L*_2_ distance and the Pearson delta (*R*^2^). (C) True positive rate (recall) of the interaction predictions as a function of the false discovery proportion. *TP*: true positive, *FP*: false positive, *FN*: false negative. (D) Classification of interactions based on the difference from the additive expectation (the error bars show the additive range). (E) Bar chart of the composition of the observed interaction classes. (F) Top: Scatterplot of observed vs. the predicted expression compared to the additive expectation. Each point is one of the 1000 read-out genes under one of the 62 double perturbations across five test-training splits. The 500 predictions that deviated most from the additive expectation are depicted with bigger and more saturated points. Bottom: Mosaic plots that compare the composition of highlighted predictions from the top panel stratified by the interaction class of the prediction. The width of the bars is scaled to match the number of instances.

Next, we considered the models’ ability to predict genetic interactions. Conceptually, a genetic interaction exists if the phenotype of two (or more) simultaneous perturbations is “surprising”, i.e., differs from the additive expectation. We used the following statistical approach to define and extract a set of interactions from the held-out data and compare it to the model predictions: For each of the 124 double perturbations and the 1 000 read-out genes, we computed the difference between the observed expression value and the additive expectation. These values showed a mixture distribution composed of a large component with a single narrow peak around zero (corresponding to a majority of non-interactions), and a smaller component consisting of two pronounced tails on either side (corresponding to interactions; Suppl. Fig. S3). To decompose this mixture, we used Efron’s empirical null approach,^14^ as implemented in the *locfdr* package. Thus, we identified a set of 5 035 interactions (out of potentially 124 000) at a false discovery rate of 5%.

For each model, we similarly computed for each of its 310 000 predictions (1 000 read-out genes, 62 held-out double perturbations across five test-training splits) the difference between prediction and additive expectation. We called a predicted interaction if the absolute value of this difference was larger than a threshold *D* > 0. We then computed for all possible choices of *D* the true positive rate (number of correctly predicted interactions divided by total number of interactions) and the false discovery proportion (number of falsely predicted interactions divided by total number of predicted interactions), which resulted in the curve shown in Fig. 1C. By definition, the additive model does not predict interactions and did not compete.

No model was better than the *no change* baseline. The same ranking of models was observed when using receiver-operating characteristic (ROC) or precision-recall curves (Suppl. Fig. S4).

To further dissect this finding, we classified the interactions using the LFCs of the observations compared to the control condition as follows (Fig. 1D), if the two individual LFCs had the same sign:

- *buffering*, if the LFC was between 0 and the additive expectation,
- *synergistic*, if it exceeded the additive expectation,
- *opposite*, if its sign differed from that of the individual perturbations.

If the individual effects were in opposite directions, *other*. According to this classification, 2.3% of the read-out gene expression values across all double perturbation were buffering interactions, 0.6% synergistic, and zero were in the opposite direction of the individual perturbations (Fig. 1E).

All models were better at identifying deviations from the additive model if the effect was buffering (Fig. 1F). We further quantified how often the models predicted synergistic effects among the 500 top interaction predictions, as this is something that neither one of our baselines could do. However, we found that the deep learning models rarely predicted synergistic effects and that it was even rarer that those predictions were correct (Fig. 1F bottom panel). To our surprise, we often found the same pair of hemoglobin genes (HBG2 and HBZ) among the top predicted interactions, across models and double perturbations (Suppl. Fig. S5). Examining the data, we noted that all models except Geneformer and scFoundation predicted LFC ≈ 0—like the *no change* baseline—for the double perturbation of these two genes, despite their strong individual effects (Suppl. Fig. S6). We confirmed that for most genes the predictions of scGPT, UCE, and scBERT did not vary across perturbations and the predictions of GEARS and scFoundation varied considerably less than the ground truth (Suppl. Fig. S7).

### Predicting the effect of unseen single perturbations

An exciting functionality of the foundation models and GEARS is their claimed ability to predict the effect of a genetic perturbation that had not previously been seen. The hope is that they have learned so much about relationships and interactions between genes (and their products) that they can extrapolate to unseen gene perturbations. For this, GEARS uses shared gene ontology^15^ annotations, while the foundation models have learned an embedding of the genes into an abstract mathematical space meant to digest all the biological phenomena they encountered in their training.

To benchmark this functionality, we used two CRISPR interference datasets by Replogle et al.^16^ obtained with K562 and RPE1 cells and a dataset by Adamson et al.^17^ obtained with K562 cells. We downloaded the data using GEARS (overview in Suppl. Fig. S1).

As a baseline, we devised a simple linear model as follows: we assume that for each read-out gene, we have a *K*_*G*_-dimensional embedding vector, and we collect these in the matrix **G** with one row per read-out gene and *K*_*G*_ columns. Similarly, for each perturbation, we have a *K*_*P*_-dimensional embedding vector, and we collect these in the matrix **P** with one row per perturbation and *K*_*P*_ columns. Given a data matrix **Y**_train_ of gene expression values, with one row per read-out gene and one column per perturbation (i.e., pseudobulked per condition of the single-cell data), the *K*_*G*_ × *K*_*P*_ matrix **W** is found as

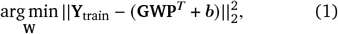

where ***b*** is a vector of the row means of the training matrix (see also schematic in Fig. 2B). To obtain embeddings **G** and **P**, we first used the following simple approach: perform a principal component analysis on **Y**_train_ and using the top *K*_*G*_ principal components for **G**. Then subset this **G** to only the rows corresponding to genes that were perturbed in the training data (and hence appear as columns in **Y**^train^) and use the resulting matrix for **P**. Having found a **W**, we can use it for prediction, 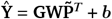, where now 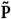 is the matrix formed by the rows of **G** corresponding to genes perturbed in the test data.

**Figure 2.**
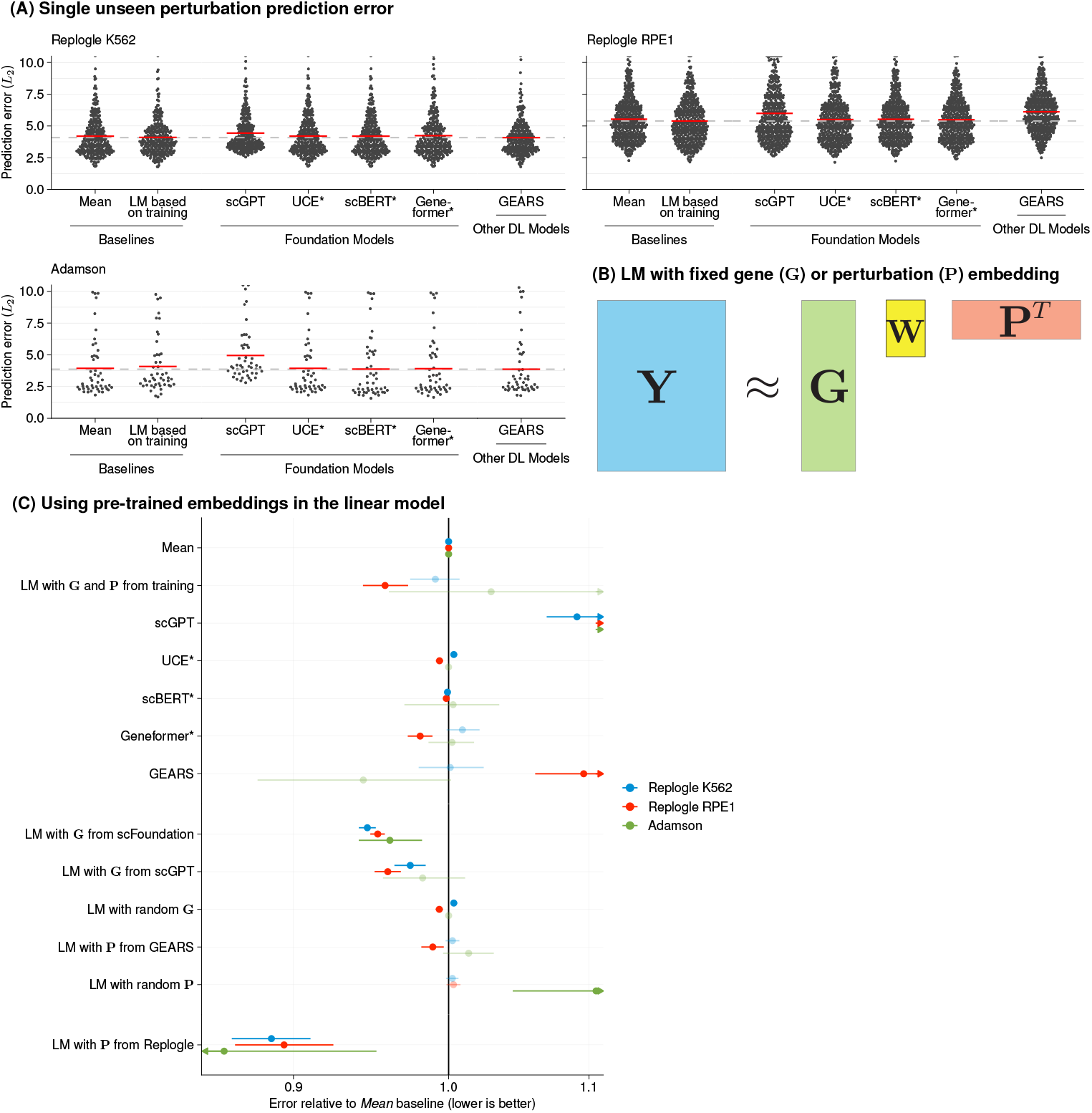
Single perturbation prediction. (A) Beeswarm plot of the prediction errors for 134, 210, and 24 unseen single perturbations across two test-training splits (Methods). The prediction error is measured by the *L*_2_ distance between the mean predicted and observed expression profile of the *n* = 1000 most highly expressed genes. The horizontal red lines show the mean per model, which for the best-performing model is extended by the dashed line. *LM*: linear model, *DL*: Deep Learning. (B) Schematic of the linear model (LM) and how it can accommodate available gene or perturbation embeddings. (C) Forest plot comparing the performance of all models relative to the error of the *mean* baseline. The point ranges show the overall mean and 95% confidence interval of the bootstrapped mean ratio between each model and the baseline. The opacity of the point range is reduced if the confidence interval contains zero.

In addition, we included an even simpler baseline, which is the mean expression across the perturbations in the training set (i.e., ***b***). This was first suggested in preprints by Kernfeld et al.^18^ and Csendes, Szalay, and Szalai^19^ that appeared while this paper was in revision.

Despite the simplicity of the mean prediction and the linear model, we found that none of the models consistently outperformed them (Fig. 2A). We obtained analogous results when using the Pearson delta measure or considering other gene sets (most expressed or most differentially expressed genes) (Suppl. Fig. S8). We did not include scFoundation in this benchmark, as it required each dataset to exactly match the genes from its pre-training data and, for the Adamson and Replogle data, the majority of the required genes were missing. We also did not include CPA, as it is not designed to predict the effects of unseen perturbations.

### Using pre-trained embeddings in a linear model

Foundation models have demonstrated the power of pretraining a model on large datasets and then fine-tuning for a specific task. To assess if the pre-training worked, we extracted the gene embeddings **G** from scFoundation and scGPT and a perturbation embedding **P** from GEARS. We find that a linear model with these embeddings performed as well or better than scGPT and GEARS with their inbuilt decoders (Fig. 2C). Furthermore, the linear models with the gene embeddings from scFoundation and scGPT outperformed the *mean* baseline and a linear model with random embeddings, but not consistently the linear model using **G** and **P** from the training data.

The approach that did consistently outperform all other models was a linear model pre-trained with **P** from Replogle (using the K562 celline data to predict the Adamson and RPE1 results, and the RPE1 cellline for the K562 results). Furthermore, we found that we could infer for which genes and perturbations the transfer would work well: genes that differed more between K562 and RPE1 were less accurately predicted (Suppl. Fig. S9). This suggests that pre-training on data that more closely matches the downstream task could boost predictive performance.

## Discussion

We presented benchmarks where current foundation models do not provide a performance benefit over deliberately simplistic linear prediction models, despite significant computational expenses for fine-tuning the deep learning models (Suppl. Fig. S10).

Our simple linear models are not intended to be serious contestants in prediction tasks, as they function without any modeling of the underlying system. Nevertheless, the foundation models benchmarked here could not surpass them. Therefore, we conclude that the goal of predicting the outcome of not-yet-performed experiments has not yet been reached. (Here, we want to reiterate that the authors of UCE, scBERT and Geneformer did not claim to be able to predict gene expression changes after perturbation, and we only included them for completeness.)

This result is not due to the rarity of interactions: we identified 5 035 read-out gene double perturbation pairs (4.2% of all observations) that did not fit the additive null model at a false discovery rate of 5%.

The publications that presented the three deep learning models each included a comparison against different baselines: the authors of GEARS compared it to a fork of CPA, scGPT was compared to GEARS and a linear model, and scFoundation was compared to GEARS. Some of these baselines may have happened to be particularly “easy”. For instance, CPA in our benchmarks appears to break down in the double perturbation benchmark and was never designed to predict the effects of unseen perturbations. The linear model used in scGPT’s benchmark appears to have been set up such that it reverts to predicting no change over the control condition for unseen perturbations.

Our results are in line with previously published benchmarks that assessed the performance of foundation models for single-cell data clustering, cell type annotation, batch correction, or perturbation prediction, which found negligible benefit compared to simpler approaches.^20,21,22^ Our results also concur with a study showing that a simple baseline can outperform GEARS for predicting unseen single perturbations^23^ and a study that compared CPA and GEARS against an additive model for double perturbation prediction.^24^ Since the release of our paper as a preprint, several other benchmarks^18,19,25,26,27,28,29,30^ were released that show that deep learning models struggle to outperform simple baselines. Two of these preprints^18,19^ suggested an even simpler model than our linear model (1), namely to simply predict the global means, and we have included their idea in our benchmark.

One limitation of our benchmark is that we only used four different datasets. We chose those as they were also used in the publications describing GEARS, scGPT, and scFoundation. Another limitation is that all datasets are from cancer cell lines, which, for example, Geneformer excluded from their training data because of concerns about their high mutational burden. We also did not exclude perturbations that did not seem to affect the level of their target gene in the data; doing so could potentially improve the quality of the training data.

We explored whether the embeddings learned by the foundational models have value independent from their inbuilt decoders and found hints that this might be the case. Furthermore, we noted that pre-training on a perturbation dataset, instead of observational data, provided a performance benefit. This leads us to hypothesize that pre-training on data that more closely matches the downstream task could improve model performance.

We do not consider our negative results on the foundation models’ performance in the prediction task as arguments against this line of research. Neural networks are well-established and have been successful in singlecell omics: e.g., autoencoder-based architectures are used widely for tasks such as denoising and interbatch and multimodal integration.^31,32^ The progress brought by the transformer architecture and the transfer learning paradigm for many machine learning tasks is real and substantial.

In the last two decades of machine learning research, including the recent successes of LLMs, reliable benchmarks with careful design and scoring were essential in guiding the field.^33^ Our work provides a benchmark for single cell transcriptome prediction tasks, together with exemplary baselines that would need to be surpassed in order to show that real, non-trivial insights have been transferred from the foundation data. We hope that this will help direct further work in bringing transfer learning to biology.

## Data Availability

All datasets used in this manuscript are publicly available:

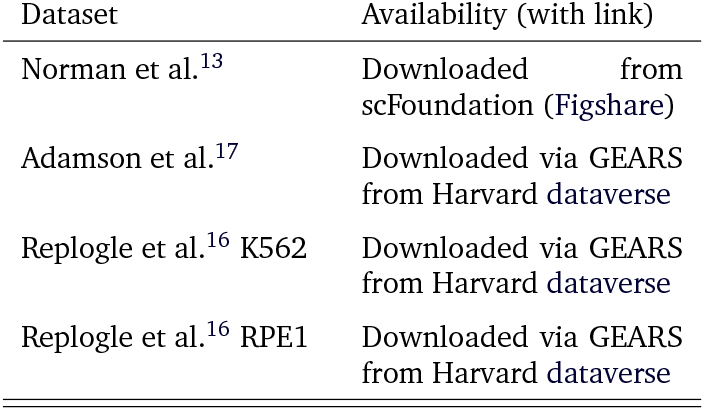

| Dataset | Availability (with link) |
| --- | --- |
| Norman et al.<sup>13</sup> | Downloaded from scFoundation (Figshare) |
| Adamson et al.<sup>17</sup> | Downloaded via GEARS from Harvard dataverse |
| Replogle et al.<sup>16</sup> K562 | Downloaded via GEARS from Harvard dataverse |
| Replogle et al.<sup>16</sup> RPE1 | Downloaded via GEARS from Harvard dataverse |

## Code Availability

The code to reproduce the analyses presented here and details about the software package versions are available at github.com/const-ae/linearperturbationprediction-Paper, which we also archived on Zenodo^34^

## Acknowledgments

We thank an anonymous reviewer of one of our previous manuscripts^35^ for the suggestion to compare foundation models against simple linear models, which eventually led to this work.

## Methods

We ran the double perturbation benchmark on the data produced by Norman et al.^13^ and reprocessed by scFoundation. For the single gene perturbation benchmarks, we used the data from Adamson et al.^17^ and Replogle et al.^16^ as provided by GEARS (details in Data Availability).

We ran GEARS version 0.1.2, scGPT version 0.2.1, scFoundation (which is build on top of a GEARS version 0.0.2 fork), CPA version 0.8.8, Geneformer version 0.1.0, scBert from commit hash 262fd4b9 with model weights provided by the authors, and UCE at commit hash 8227a65c. We used each model, as much as possible, with their default parameters. All scripts that were used to predict the expression changes are available on GitHub (github.com/const-ae/linearperturbationprediction-Paper/tree/main/benchmark/src).

- GEARS and scFoundation provide a straightforward API to predict the expression change after perturbation. We limited the fine-tuning time to three days, which meant that we trained scFoundation for five epochs.
- For scGPT, we used the same parameters and code as in their tutorial for perturbation prediction.
- For CPA, we used the code from their tutorial on how to predict combinatorial CRISPR perturbations on the Norman dataset.
- For Geneformer, we fine-tuned the provided model by predicting the perturbation labels of the training data. We then used the built-in *in silico* perturbation functionality to calculate the perturbed embedding.
- UCE is designed for zero-shot use, which means that it does not need to be fine-tuned. We report results from the four-layer version of UCE (as we found no performance difference between the four or 33-layer versions). UCE does not provide functionality for *in silico* perturbation, so we calculated the post-perturbation embedding by taking the expression matrix for the unperturbed cells and overwrote the rows for the genes that we wanted to perturb with the values from the ground truth expression matrix. We thus tried to ensure that we tested the model under the best conditions, accepting that test data leakage could theoretically give the model an advantage over the other models.
- We fine-tuned scBERT on predicting the perturbation labels of the training data. We then used the same approach to calculate the embedding after *in silico* perturbation that we used for UCE.

To predict the expression changes from the embeddings of Geneformer, UCE, and scBERT, we added a linear decoder to the models. We fitted a ridge regression model that predicted the gene expression of the perturbed cells from the perturbed embeddings of the training data. We then used that ridge regression to predict the gene expression of the test data from the corresponding perturbed embeddings and continued with the mean of the predicted values per perturbation.

To reduce the probability that we understate the performance for any of the models, due to wrong or suboptimal operation by ourselves, we reached out to the original authors of the benchmarked models and asked them to review our code. The authors of CPA perceived a problem with our code and submitted a fix^1^; however, as the new code had worse performance than the original version, we here report results of the original code.

For the double perturbation benchmark, we split the data into test and training sets. We assigned all single gene perturbations and a randomly chosen half of the double perturbations to the training and used the other half of the double perturbations as the test set. To reduce stochastic effects on our results, we repeated the whole procedure including the random test-training splitting five times. For the single perturbation benchmark, we used GEARS’ *simulation* test-training splitting procedure, which we repeated twice.

For the double perturbation benchmark, we used two baseline models: no change and additive. The *no change* model “predicted” for each double perturbation the expression values seen in the control condition (***y***^∅^). The *additive* model predicts the expression after a double perturbation of genes A and B as

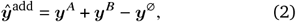

where ***y*** ^*A*^ and ***y***^*B*^ are the mean observed expression vectors for the single perturbation of gene A and B, respectively.

To predict the effects of unseen single perturbations, we used two baselines. The *mean* model calculated the mean of the expression values in the training data. The *linear model* is implied in Eqn. (1). We set ***b*** to the row means of the training data 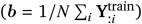 and find **W** using the normal equations

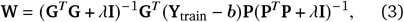

where we use a ridge penalty of λ = 0.1 for numerical stability.

For the single perturbation analysis, not all models were able to predict the expression change for all unseen perturbations. For example, the linear model with **G** and 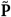 from the training data could only predict perturbations where the target genes was also part of the read-out genes. To evaluate all models on a consistent set of perturbations, we restricted our analysis to those perturbations for which we had predictions from all models (73 perturbations for Adamson, 398 for Replogle K562, and 629 for Replogle RPE1).

We converted GEARS’ gene ontology annotations into a perturbation embedding **P** by computing a spectral embedding^36,37^ of pathway membership matrix. We extracted the gene embedding **G** from scGPT following their tutorial on gene regulatory inference. For scFoundation, we extract **G** directly from the pre-trained model weights (*pos emb*.*weight*). For the linear model with **P** from the Replogle data, we fitted a ten-dimensional PCA on the columns of the matrix with the perturbation means of the reference data. We fitted all linear models as described in the main text with *K*_*G*_ = 10; if **G** or **P** were provided, we simply replaced the estimate from the training data with the provided matrix before calculating **W**.

The additive model is a special case of the linear model (1) where the gene embedding is simply the single perturbation data, without any further transformation or dimension reduction (**G** = **Y**^single^), the perturbation embedding **P** is a binary coding, where each column vector has ones in the rows of the perturbed genes and is zero otherwise, and *W* is an identity matrix and ***b*** = −***y***^∅^.

We measured the prediction error using the distance 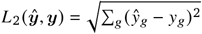 (also called root mean squared error) between the observed expression values and predictions for the 1 000 most highly expressed genes in the control condition. We also calculated the Pearson delta correlation metric, as suggested by Cui et al.^9^: PearsonDelta (*ŷ, y*) = cor(*ŷ*−*y*^∅^, *y*−*y*^∅^). Unlike the *L*_2_ distance, the Pearson delta metric does not penalize predictions that are consistently too small or too large in amplitude, and thus prioritizes correct prediction of the direction of the expression change.

For the double perturbation data, we assess the true positive rate (recall) as a function of the false discovery rate. First, we find the order statistic of absolute difference of predictions and additive expectation across all test perturbations (*j* = argsort(vec(|Ŷ−Ŷ^add^|))), where Ŷ is the matrix of the predictions for all genes and perturbations and Ŷ^add^ are the additive expectations.

The false discovery proportion (FDP) at position *l* ∈ {1, …, *N*} for a threshold *u*, that separates the interactions from the non-interactions, is

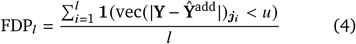

and the true positive rate (TPR) is

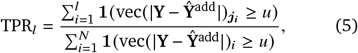

where **Y** is the matrix of observed value and *N* is the product of the number of genes and perturbations. The indicator function **1**(·) counts how often the observed values **Y** deviate enough from the additive expectation so that the observations is considered an interaction. The order statistic ***j*** ensures we consider the gene-perturbation pairs first, where the model prediction deviates most from the additive expectation.

Lastly, we find the order statistic of the false discovery proportions (***s*** = argsort(FDP)) and plot the tuples 1, …, *N*

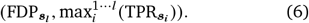

The advantage of the false discovery vs. true positive curve over the traditional precision-recall or receiver operator curve is that it provides a direct assessment which fraction of interactions a model identifies for a fixed fraction of false positives.

## Supplement

**Suppl. Figure S1.**
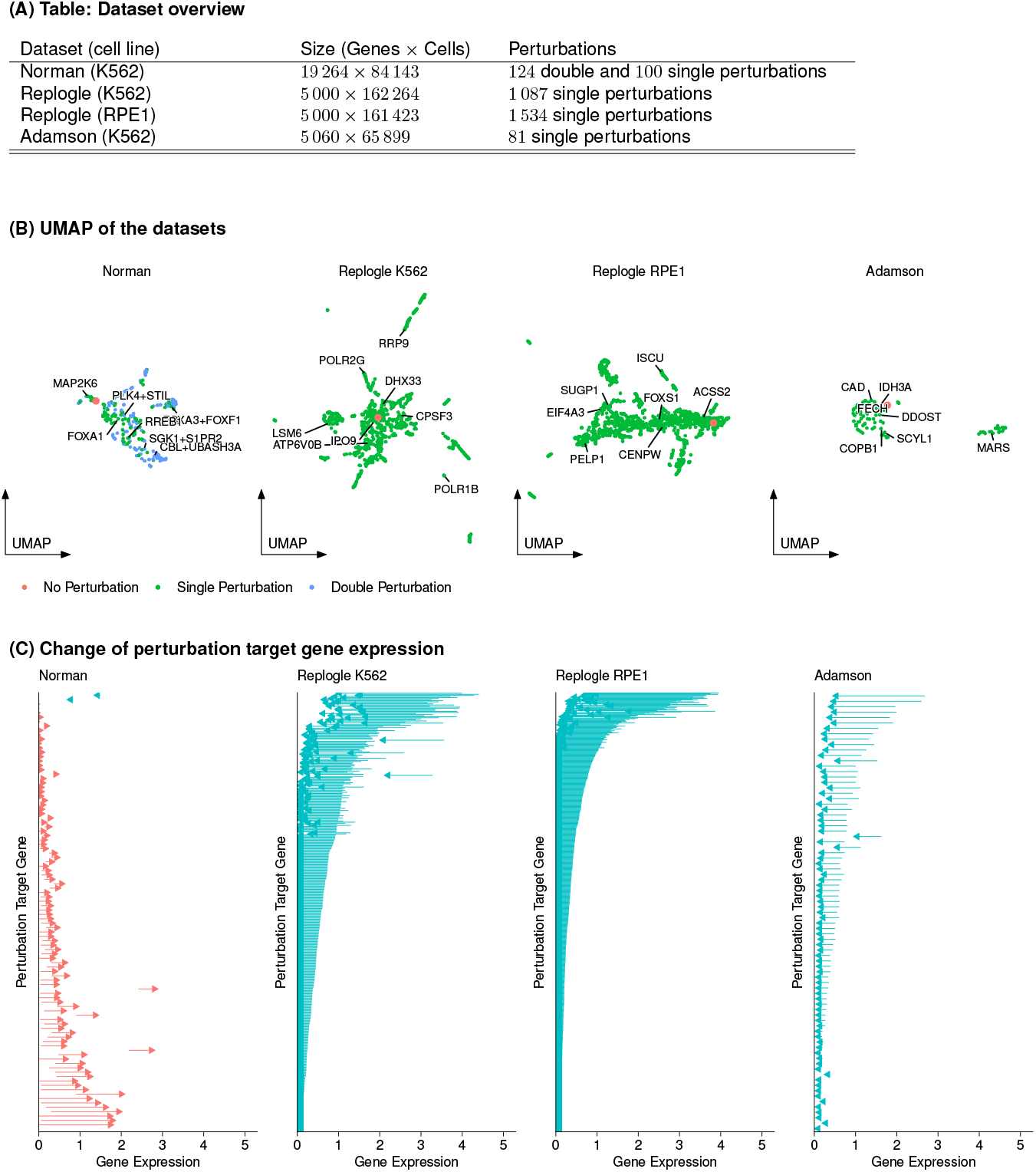
Dataset Overview. (A) Table with the size of the data and the number of perturbations. (B) UMAP on the perturbations per dataset (aggregated to the mean per perturbation). The position of the control condition without perturbation is shown in red, and a random selection of perturbations is labeled. (C) Change in the expression of the target gene of each perturbation. The base of the arrow indicates the expression without perturbation, and the tip indicates the expression after perturbation. For genes targeted multiple times in the Norman dataset, we show the average expression after perturbation.

**Suppl. Figure S2.**
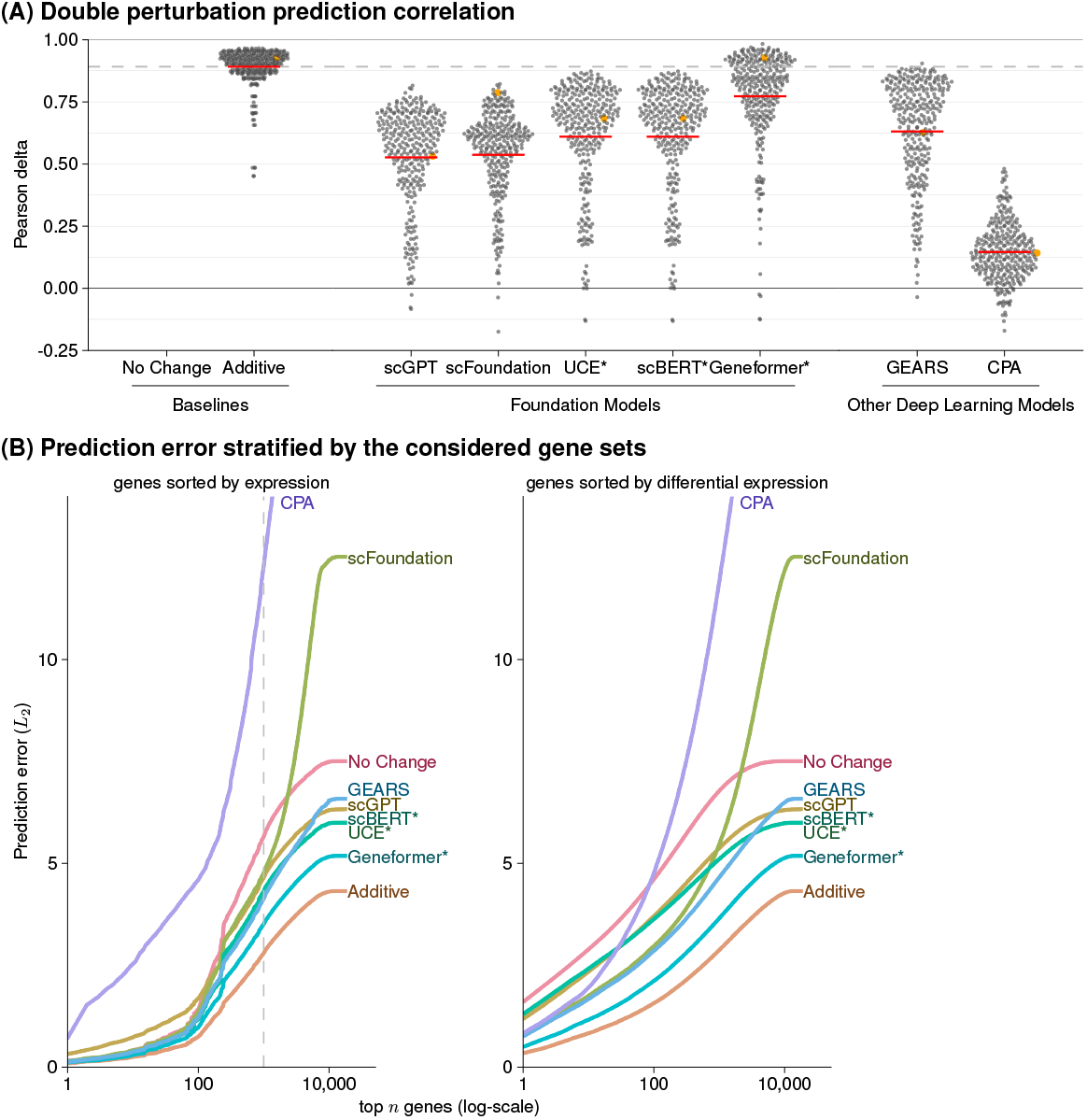
Alternative measures of the double perturbation prediction performance. (A) The Pearson delta measure calculates the correlation of the prediction and observations after subtracting the expression in the control condition. The correlation for the *no change* predictions could not be calculated because they were all zero. The horizontal red lines show the mean per model, and the dashed line indicates the correlation of the best-performing model. (B) Prediction error as a function of *n*, the number of read-out genes. Left: genes ranked by expression in the control condition, right: by differential expression between observed value and expression in the control condition. Note that sorting by differential expression is only possible if access to the ground truth is available and can thus not be applied in real-world use cases. The dashed line at *n* = 1000 is the choice in Panel A and elsewhere in this work.

**Suppl. Figure S3.**
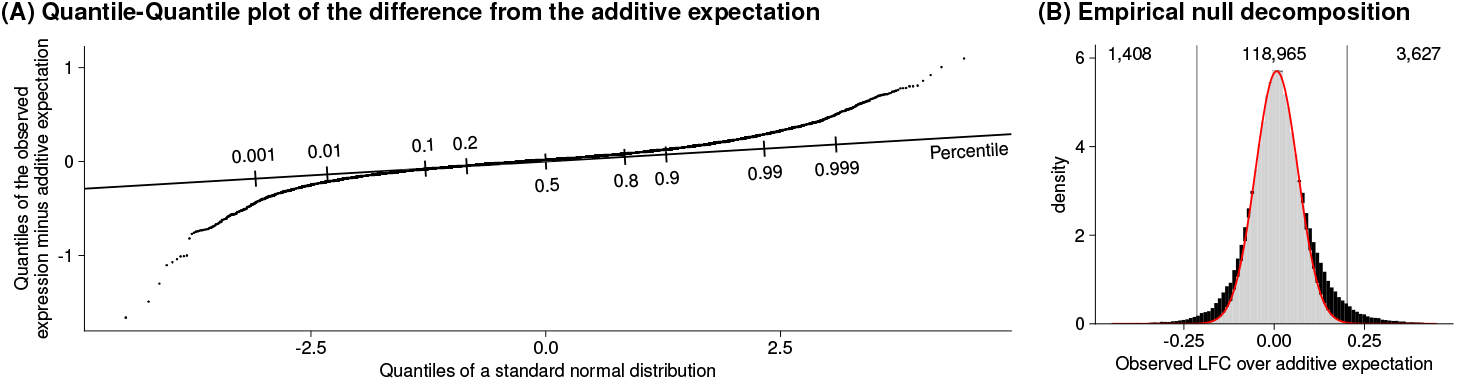
Distribution of the observed difference from the additive model. (A) Quantile-quantile plot comparing the distribution of the differences between observed expression values and the additive expectation against a standard normal distribution. The slope of the line is the standard deviation of the null model. (B) Histogram of the differences with a red curve overlayed that shows the null distribution fitted using *locfdr*. Values under the curve are grey, and the black bars show the observations that exceed what we would expect under the null model. The vertical bar shows the upper and lower thresholds for which the observations have a false discovery rate of less than 5% (i.e., the grey fraction of the bars outside the vertical lines is 5%). The numbers at the top count the observations per group.

**Suppl. Figure S4.**
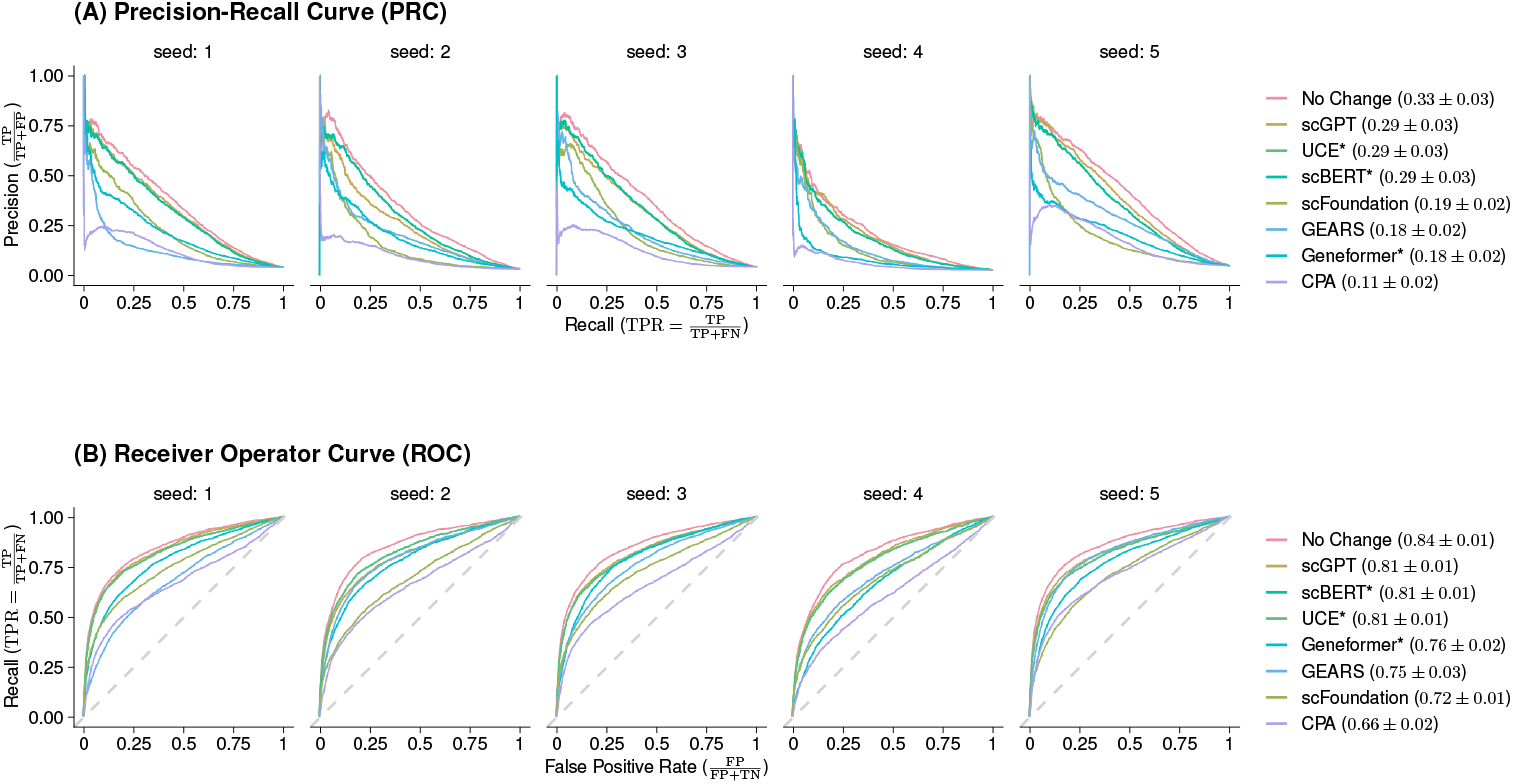
Alternative measures how well each model detects genetic interactions. (A) Precision-recall and (B) receiver operator curve for all models distinguishing interactions from additive combinations. The numbers in parenthesis are the area under the curve (AUC) with the standard error across five test-training splits. *TP*: true positive, *FP*: false positive, *FN*: false negative, *TN*: true negative.

**Suppl. Figure S5.**
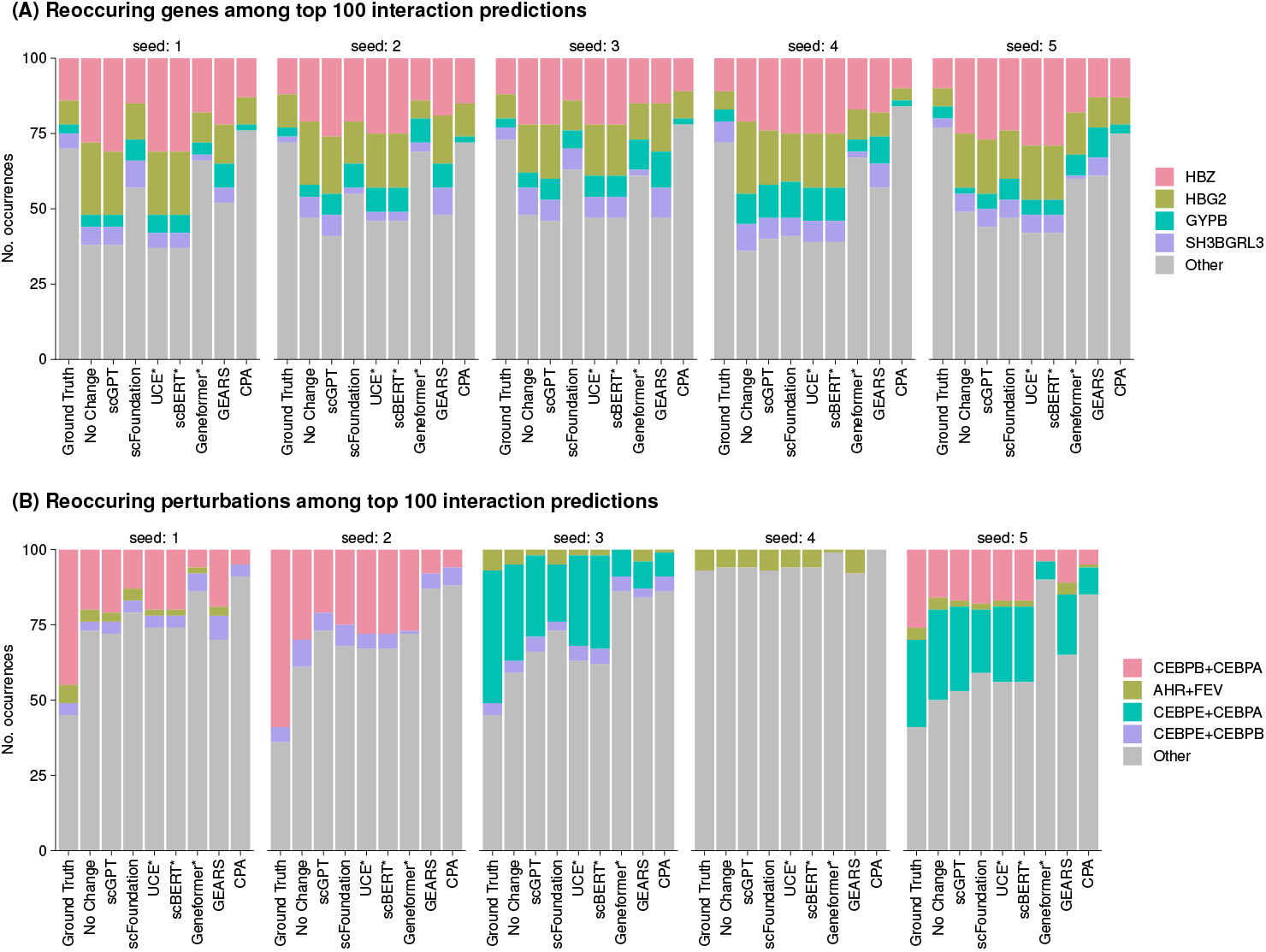
Reoccurrence of genes and perturbations for top predictions. (A) Reoccurrence of genes and (B) perturbations among the 100 predictions that differed most from the additive expectation. The data is facetted by the test-training split. The ground truth column shows the genes and perturbations sorted by observed difference from the additive expectation. The highlighted genes and perturbations are the six most reoccurring ones.

**Suppl. Figure S6.**
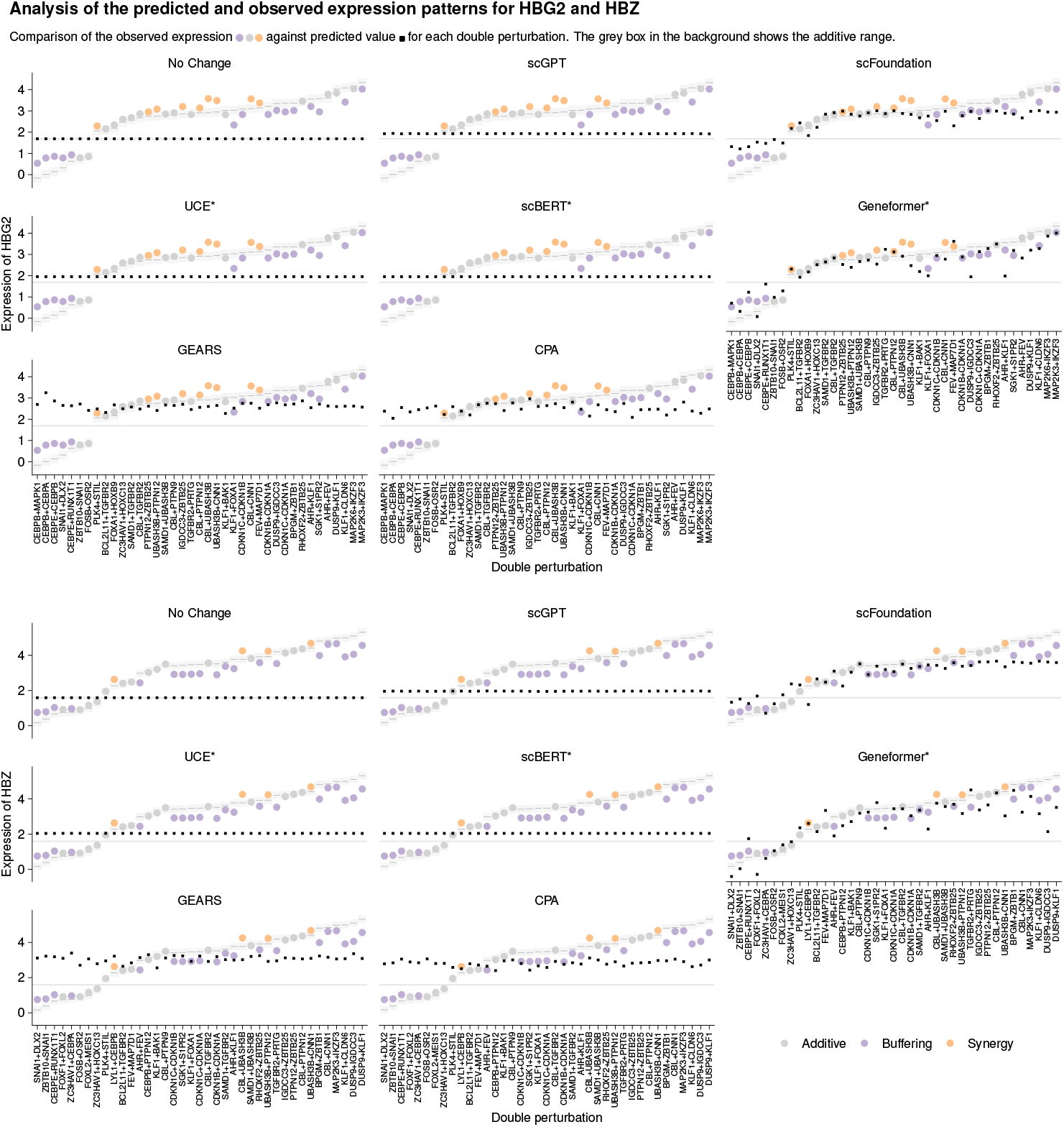
Comparison of predicted and observed expression for HBG2 and HBZ. Comparison of the predicted expression (black squares), the observed expression values (points colored by interaction type), and the range of values that are considered additive (grey boxes) for all test perturbations with seed = 1. The grey horizontal line shows the expression of HBG2 and HBZ without perturbation.

**Suppl. Figure S7.**
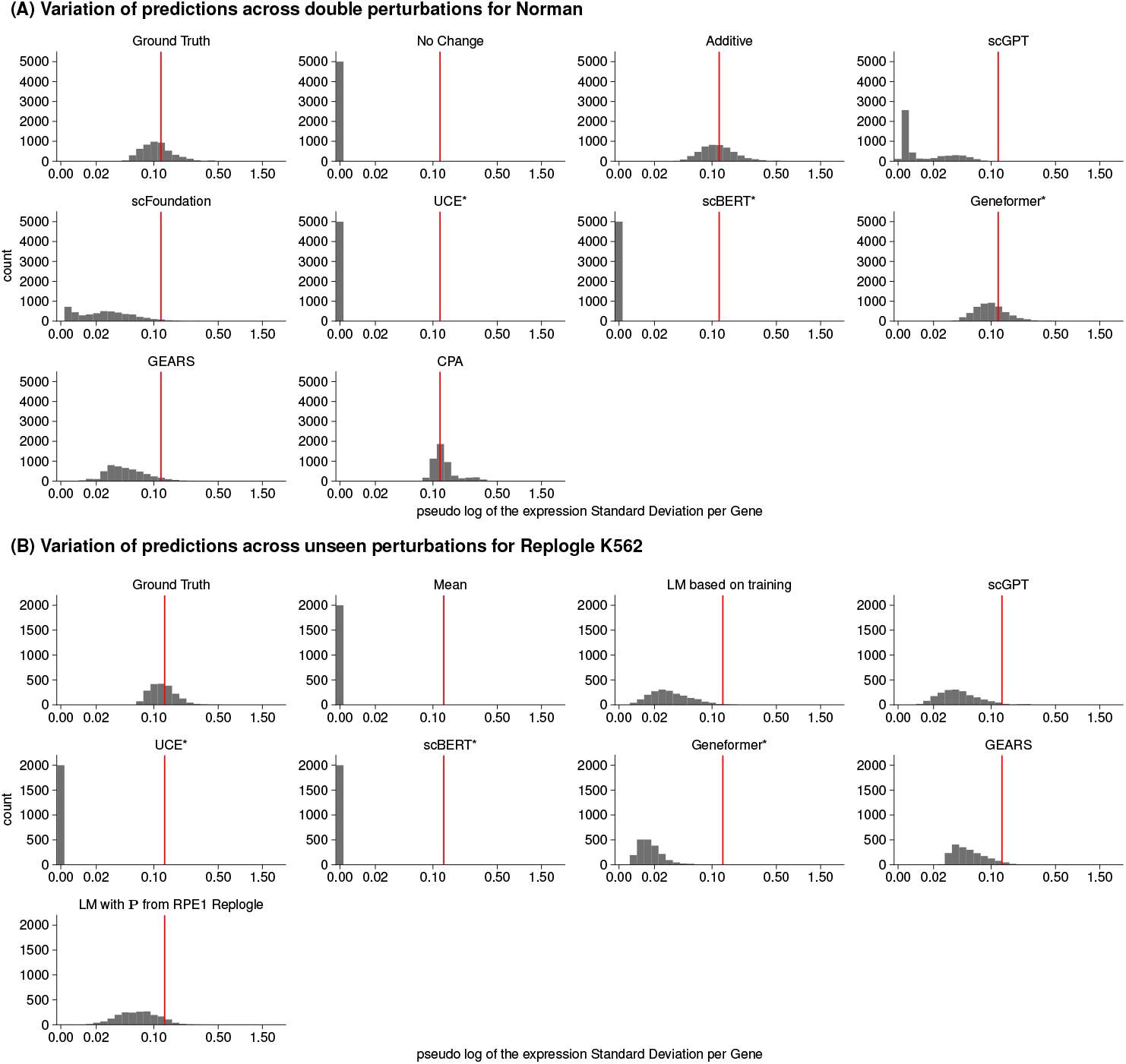
Variation of the predicted and observed expression values. Histogram of the standard deviation per gene for the predicted and observed expression values across perturbations facetted by the model. The red vertical bar indicates the mean of the standard deviations for the ground truth for (A) the Norman dataset and (B) the Replogle K562 dataset. The data reflects the variation for the 1 000 most highly expressed genes and is aggregated across five test-training splits. *LM*: linear model.

**Suppl. Figure S8.**
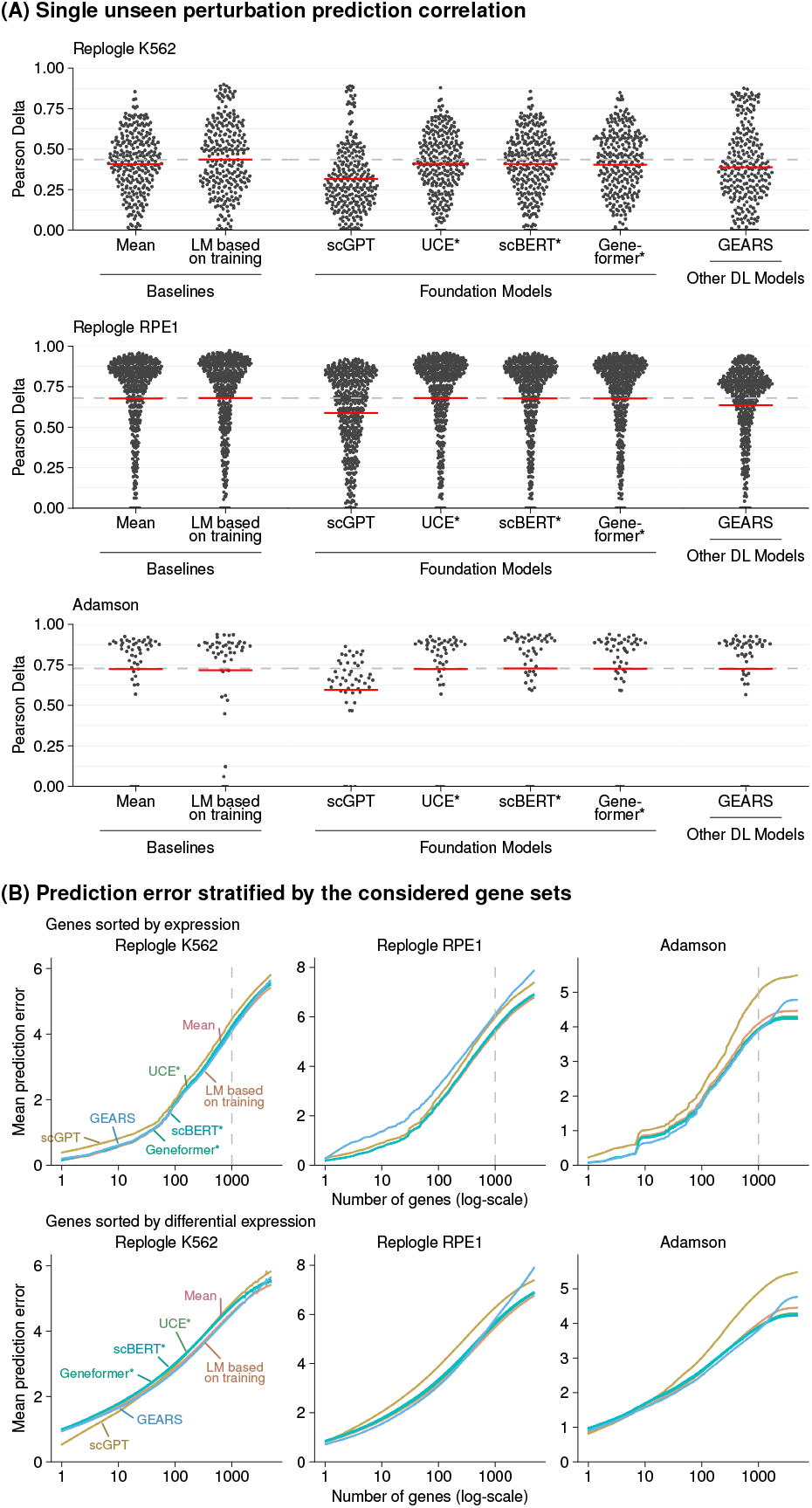
Alternative measures of the single perturbation prediction performance. (A) The Pearson delta measure calculates the correlation of the prediction and observations after subtracting the expression in the control condition. The horizontal red lines show the mean per model and the dashed line indicates the correlation of the best-performing model. (B) Prediction error as a function of *n*, the number of read-out genes. Top: genes ranked by expression in the control condition, bottom: by differential expression between observed value and expression in the control condition. Note that sorting by differential expression is only possible if access to the ground truth is available and can thus not be applied in real-world use cases. The dashed line at *n* = 1000 is the choice in Panel A and elsewhere in this work. *LM*: linear model, *DL*: deep learning.

**Suppl. Figure S9.**
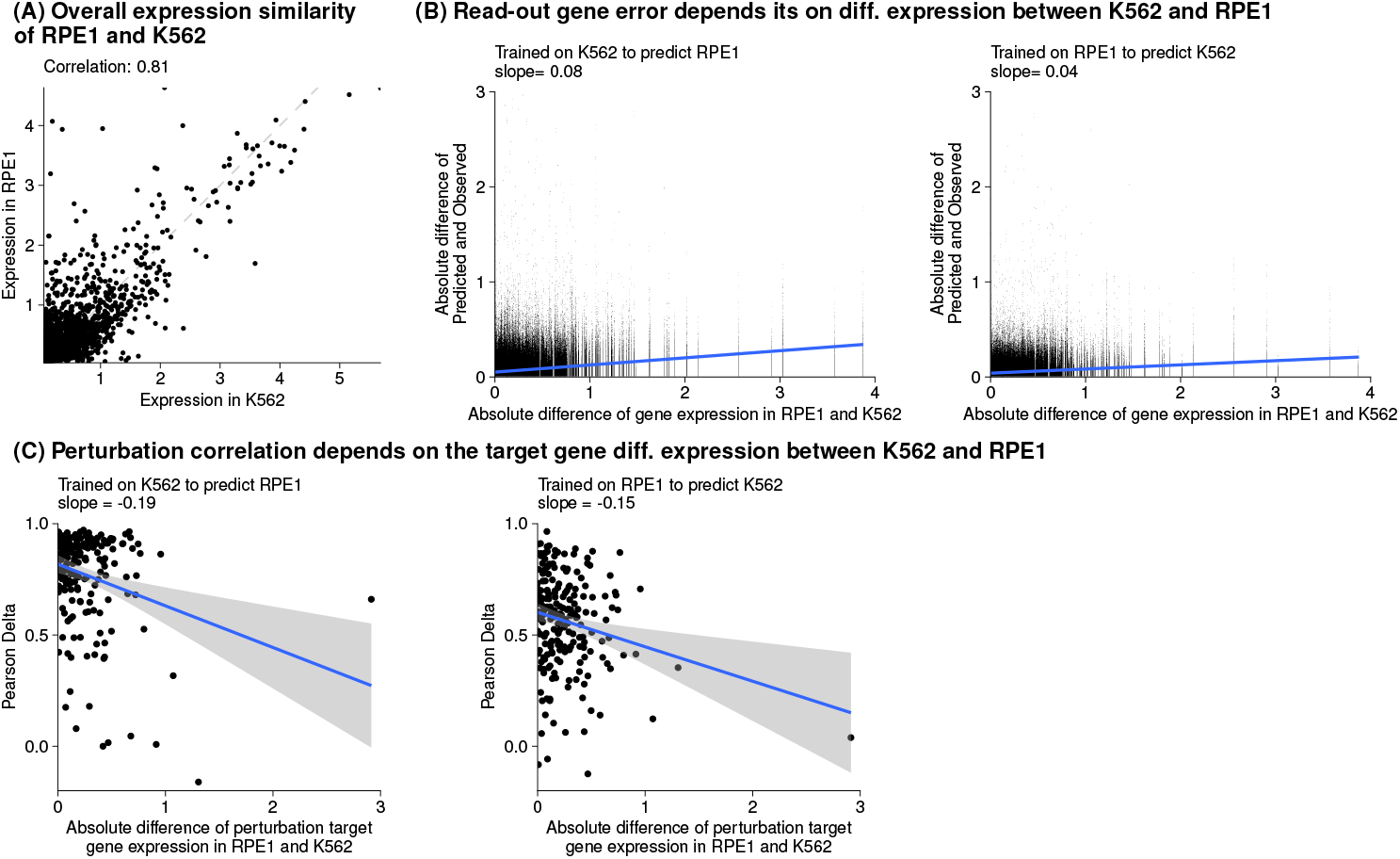
Analysis how differential expression between K562 and RPE1 effects prediction accuracy of transfer learning. (A) Scatter plot of the mean gene expression for shared genes between RPE1 and K562 without perturbation. The dashed line indicates the diagonal. (B) Scatter plot of the absolute prediction error per read-out gene against the differential expression of that gene between RPE1 and K562. Each point is one read-out gene from one of the 122 double perturbations from five test-training splits. The blue line shows the linear fit with a slope indicated in the subtitle. (C) Scatter plot of the Pearson delta score per perturbation for the RPE1 dataset against the differential expression of the perturbation target gene between RPE1 and K562. The blue line shows the linear fit with a slope indicated in the subtitle, and the shaded area indicates the standard error of the fit.

**Suppl. Figure S10.**
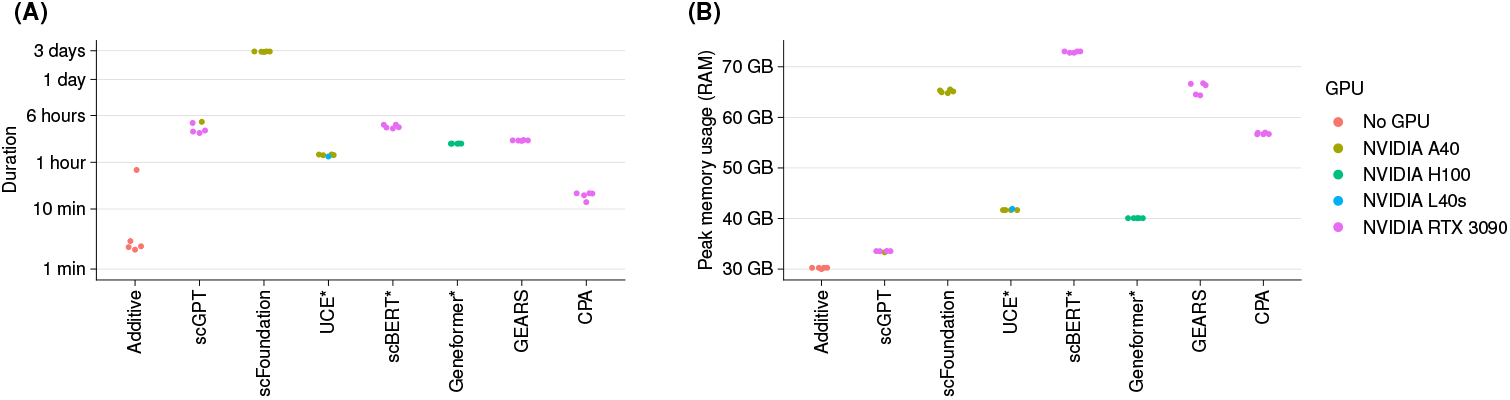
Computational resource requirements. The resource usage was measured for the Norman dataset, which had 19 624 genes and 81 143 cells grouped into 225 conditions. Each point is one of the five test-training splits. (A) Elapsed time on a log scale to fine-tune and predict the double perturbations. (B) Peak memory usage for each model was measured using GNU time. The points are colored by the respective GPU model that was used.

## Footnotes

1 https://github.com/const-ae/linear_perturbation_prediction-Paper/pull/3

## References

[1] George I Gavriilidis et al. “A mini-review on perturbation modelling across single-cell omic modalities”. In: Computational and Structural Biotechnology Journal (2024). DOI: 10.1016/j.csbj.2024.04.058.

[2] Artur Szałata et al. “Transformers in single-cell omics: a review and new perspectives”. In: Nature Methods 21.8 (2024), pp. 1430–1443. DOI: 10.1038/s41592-024-02353-z.

[3] Jennifer E. Rood, Anna Hupalowska, and Aviv Regev. “Toward a foundation model of causal cell and tissue biology with a Perturbation Cell and Tissue Atlas”. In: Cell 187.17 (Aug. 2024), 4520–4545. ISSN: 0092-8674. DOI: 10.1016/j.cell.2024.07.035.

[4] Aviv Regev et al. “The human cell atlas”. In: eLife 6 (2017), e27041. DOI: 10.7554/eLife.27041.

[5] CZI Single-Cell Biology et al. “CZ CELLxGENE Discover: A single-cell data platform for scalable exploration, analysis and modeling of aggregated data”. In: bioRxiv (2023), pp. 2023–10. DOI: 10.1101/2023.10.30.563174.

[6] Fan Yang et al. “scBERT as a large-scale pretrained deep language model for cell type annotation of single-cell RNA-seq data”. In: Nature Machine Intelligence 4.10 (2022), pp. 852–866. DOI: 10.1038/s42256-022-00534-z.

[7] Christina V Theodoris et al. “Transfer learning enables predictions in network biology”. In: Nature 618.7965 (2023), pp. 616–624. DOI: 10.1038/s41586-023-06139-9.

[8] Yanay Rosen et al. “Universal cell embeddings: A foundation model for cell biology”. In: bioRxiv (2023), pp. 2023–11. DOI: 10.1101/2023.11.28.568918.

[9] Haotian Cui et al. “scGPT: toward building a foundation model for single-cell multi-omics using generative AI”. In: Nature Methods (2024), pp. 1–11. DOI: 10.1038/s41592-024-02201-0.

[10] Minsheng Hao et al. “Large-scale foundation model on single-cell transcriptomics”. In: Nature Methods (2024), pp. 1–11. DOI: 10.1038/s41592-024-02305-7.

[11] Yusuf Roohani, Kexin Huang, and Jure Leskovec. “Predicting transcriptional outcomes of novel multigene perturbations with GEARS”. In: Nature Biotechnology 42.6 (2024), pp. 927–935. DOI: 10.1038/s41587-023-01905-6.

[12] Mohammad Lotfollahi et al. “Predicting cellular responses to complex perturbations in high-throughput screens”. In: Molecular Systems Biology (2023), e11517. DOI: 10.15252/msb.202211517.

[13] Thomas M Norman et al. “Exploring genetic interaction manifolds constructed from rich single-cell phenotypes”. In: Science 365.6455 (2019), pp. 786–793. DOI: 10.1126/science.aax4438.

[14] Bradley Efron. Large-scale inference: empirical Bayes methods for estimation, testing, and prediction. Vol. 1. Cambridge University Press, 2012. DOI: 10.1017/CBO9780511761362.

[15] The Gene Ontology Consortium. “The Gene Ontology knowledgebase in 2023”. In: Genetics 224.1 (2023). DOI: 10.1093/genetics/iyad031.

[16] Joseph M Replogle et al. “Mapping information-rich genotype-phenotype landscapes with genome-scale Perturb-seq”. In: Cell 185.14 (2022), pp. 2559–2575. DOI: 10.1016/j.cell.2022.05.013.

[17] Britt Adamson et al. “A multiplexed single-cell CRISPR screening platform enables systematic dissection of the unfolded protein response”. In: Cell 167.7 (2016), pp. 1867–1882. DOI: 10.1016/j.cell.2016.11.048.

[18] Eric Kernfeld et al. “A systematic comparison of computational methods for expression forecasting”. In: bioRxiv (2024). DOI: 10.1101/2023.07.28.551039.

[19] Gerold Csendes, Kristóf Z Szalay, and Bence Szalai. “Benchmarking a foundational cell model for post-perturbation RNAseq prediction”. In: bioRxiv (2024), pp. 2024–09. DOI: 10.1101/2024.09.30.615843.

[20] Kasia Zofia Kedzierska et al. “Assessing the limits of zero-shot foundation models in single-cell biology”. In: bioRxiv (2023), pp. 2023–10. DOI: 10.1101/2023.10.16.561085.

[21] Rebecca Boiarsky et al. “A deep dive into single-cell RNA sequencing foundation models”. In: bioRxiv (2023), pp. 2023–10. DOI: 10.1101/2023.10.19.563100.

[22] Tianyu Liu et al. “Evaluating the Utilities of Foundation Models in Single-cell Data Analysis”. In: bioRxiv (2024). DOI: 10.1101/2023.09.08.555192.

[23] Kaspar Märtens, Rory Donovan-Maiye, and Jesper Ferkinghoff-Borg. “Enhancing generative perturbation models with LLM-informed gene embeddings”. In: ICLR 2024 Workshop on Machine Learning for Genomics Explorations. 2024. URL: https://openreview.net/forum?id=eb3ndUlkt4.

[24] Thomas Gaudelet et al. “Season combinatorial intervention predictions with Salt & Peper”. In: ICLR 2024 Workshop on Machine Learning for Genomics Explorations. 2024. URL: https://openreview.net/forum?id=Wj95feICkN.

[25] A. Wenteler et al. “PertEval-scFM: Benchmarking Single-Cell Foundation Models for Perturbation Effect Prediction”. In: bioRxiv (2024). DOI: 10.1101/2024.10.02.616248.

[26] Ihab Bendidi et al. Benchmarking Transcriptomics Foundation Models for Perturbation Analysis : one PCA still rules them all. 2024. DOI: 10.48550/arXiv.2410.13956.

[27] Yan Wu et al. PerturBench: Benchmarking Machine Learning Models for Cellular Perturbation Analysis. Nov. 2024. DOI: 10.48550/arXiv.2408.10609.

[28] Lanxiang Li et al. “A Systematic Comparison of Single-Cell Perturbation Response Prediction Models”. In: bioRxiv (2024). DOI: 10.1101/2024.12.23.630036.

[29] Chen Li et al. “Benchmarking AI Models for In Silico Gene Perturbation of Cells”. In: bioRxiv (2025). DOI: 10.1101/2024.12.20.629581.

[30] Daniel R. Wong, Abby S. Hill, and Robert Moccia. “Simple controls exceed best deep learning algorithms and reveal foundation model effectiveness for predicting genetic perturbations”. In: bioRxiv (2025). DOI: 10.1101/2025.01.06.631555.

[31] Adam Gayoso et al. “A Python library for probabilistic analysis of single-cell omics data”. In: Nature Biotechnology (2022). DOI: 10.1038/s41587-021-01206-w.

[32] Malte D Luecken et al. “Benchmarking atlas-level data integration in single-cell genomics”. In: Nature Methods 19.1 (2022), pp. 41–50. DOI:10.1038/s41592-021-01336-8.

[33] Yupeng Chang et al. “A Survey on Evaluation of Large Language Models”. In: ACM Transactions on Intelligent Systems and Technology 15.3 (2024). DOI: 10.1145/3641289.

[34] Constantin Ahlmann-Eltze, Wolfgang Huber, and Simon Anders. Zenodo repository with the code to reproduce our analyses. 2025. DOI: 10.5281/zenodo.14832393.

[35] Constantin Ahlmann-Eltze and Wolfgang Huber. “Analysis of multi-condition single-cell data with latent embedding multivariate regression”. In: Nature Genetics (2025). DOI: 10.1038/s41588-024-01996-0.

[36] Daniel L Sussman et al. “A consistent adjacency spectral embedding for stochastic blockmodel graphs”. In: Journal of the American Statistical Association 107.499 (2012), pp. 1119–1128. DOI: 10.1080/01621459.2012.699795.

[37] Gabor Csardi and Tamas Nepusz. “The igraph software package for complex network research”. In: InterJournal Complex Systems (2006), p. 1695. URL: https://igraph.org.

